## Supplemental legends for "Early and Delayed STAT1-Dependent Responses Drive Local Trained Immunity of Macrophages in the Spleen"

**Supplementary Legends**

**Figure 1 – figure supplement 1. Single Cell Gating Strategy and Populations-Specific DEGs** (**A**) Flow cytometry gating schema for identifying MPs within the splenic myeloid subset. Gating and populations are representative of a BCG inoculated mouse two weeks post injection. Lineage (Lin) staining combines CD19 and CD3e for separation of B and T cells. (**B**) DEGs between PBS and BCG vaccinated mice for the sorted NCM, mcDC2, and NK populations.

**Figure 2 – figure supplement 1.** **Kinetics Gating Strategy and Marker and Gene Expression Across Time and Populations** (**A**) Flow cytometry gating schema for identifying MPs and TI-associated markers (including CXCL9^+^ CMs [CM-Ts] and Sca-1) within the splenic myeloid subset. Gating and populations are representative of a BCG inoculated mouse two weeks post injection. Lineage staining combines CD19, CD3e, Ly6g, and NK1.1 for separation of B, T, neutrophil, and NK cells respectively. (**B-C**) Percentage of MP populations (**B**) and frequency of CM-Ts and SCA-1+ cells (**C**) from PBS (gray) and BCG (black) mice. CM, NCM, and dendritic cell ratio calculated from CD11b+ population. RPM ratio calculated from Lin- population (control: n=2-3, BCG: n=4). (**D**) Frequency of CM-Ts and SCA-1+ cells five days post PBS (gray) or BCG (black) vaccination (control: 3, BCG = 5). (**E**) Frequency of CM-Ts and SCA-1+ NCMs twenty-four hours post *S*.Tm challenge at two or eight weeks after PBS (gray) or BCG (black) vaccination (PBS: n=6, BCG: n=6). (**F**) PCA projection of all bulk splenic conditions onto the space of the two leading principal components based on all expressed genes due to training. Conditions include each time point (control: n=2-3, BCG: n=3-4). Shade intensity represents time point, diamond (PBS) or circle (BCG). (**G**) Log2 expression of selected training-associated genes over time (control: n=2-3, BCG: n=3-4). (**H**) Experimental setup for tracking expression of training markers inherited from trained BM progenitors. After initial inoculation of PBS-ip (n=2) or BCG-ip (n=2), mice were sacrificed, and BM extracted and mixed for each pair of donors. (**I**) Flow cytometry gating schema for identifying MPs and TI-associated markers (CXCL9 and Sca-1) across CD45.1/CD45.2 donors. (**J**) NCM fraction expressing Sca-1 from cells derived from control (CD45.2) or BCG (CD45.2) mice (n=7). Square or triangle represent mice who received combined BM from donor set one or two. Significance between control (CD45.2) and BCG (CD45.1) is indicated. Data in **B-E** are presented as mean±SEM. For line graphs **B** and **C**, significance represents each time point against PBS at day 3. Two-tailed *t*-test used for data in **B-E,** paired t-test for **J (***P≥0.05, **P≥0.01, ***P≥0.005, ****P≥0.001**)**.

**Figure 3 – figure supplement 1. Lineage Tracing Gating Strategy and Origin-Specific DEGs** (**A**) Flow cytometry gating schema for identification of local and recruited MPs in the spleen. Gating and populations are representative of a BCG inoculated mouse two weeks post injection. (**B**) Percent of splenic monocytes double positive for CX3CR1 and TdTomato between PBS (gray) and BCG (black) vaccinated mice (PBS: n=4, BCG: n=5). (**C**) DEGs between PBS and BCG vaccinated mice for the bulk sorted CM, NCM, and Tdtm+/- RPM. (**D**) Expression of genes enriched within the IFNy response term across Tdtm+/- RPM. Data in **B** are presented as mean±SEM.

**Figure 4 – figure supplement 1. Training Inhibition with Stat-1 KO Mice and Across Interferon Inhibition Strategies.** (**A**) Mouse model to detect training markers in BCG-vaccinated WT and STAT1-KO mice. (**B**) CXCL9 and Sca-1 expression in MPs between control and BCG mice in WT or STAT1-KO background. (**C**) CD11b^+^ and RPM populations between control and BCG mice in WT or STAT1-KO background. (**D**) Mouse model of BCG vaccination with early interferon inhibition using the small molecule inhibitor Fedratinib targeting Type-II IFN, or Deucravacitinib targeting Type-I IFN. (**E**) Heatmap of log2 fold change for DEGs across inhibitor conditions. (**F**) CXCL9 expression in MPs across inhibitor conditions. (**G**) Flow cytometry gating strategy used for identification of BM LSK HSCs. Gating is representative of a BCG inoculated mouse two weeks post injection. Lineage staining combines CD4, CD8, B220, Ter119, Gr1, B220, and CD11b for separation of T, B, erythroid, granulocyte, and myeloid cells respectively. (**H**) Flow cytometry plot of LSK expansion within the BM across inhibitor conditions. (**I**) Two-dimensional spleen area from across experimental conditions including training, Fedratinib inhibitor, and *S.Tm* infection (n=3-6). Significance between inhibitor and infection conditions is indicated. (**J**) PCA projection of all samples onto the space of the two leading principal components based on all DEGs across training, inhibitor, and infection conditions (two-sample t-test, 5% FDR). The percentage of variance explained by each PC is indicated at the PC axes. Color is indicative of control or BCG mice before and after *S.Tm* challenge. Circle or diamond shape represent treatment with DMSO or Fedratinib. (**K-L**) Heatmap of STAT-1 signature genes (**K**) and their mean log2 expression across training conditions and antibody treatment (**L**). (**M**) Total BCG CFU from spleens from isotype and α-IFNγ treated mice two weeks post vaccination (n=4). Data in **I** and **M** are presented as mean±SEM. Heatmap rows in **E** and **K** indicate biological replicates. Two-tailed *t*-test used for data in **I** and **L (***P≥0.05, **P≥0.01, ***P≥0.005, ****P≥0.001**)**.
