## Supplementary figures and images for "Early and Delayed STAT1-Dependent Responses Drive Local Trained Immunity of Macrophages in the Spleen"

### Supplement Figure 1

**Figure supplement 1**

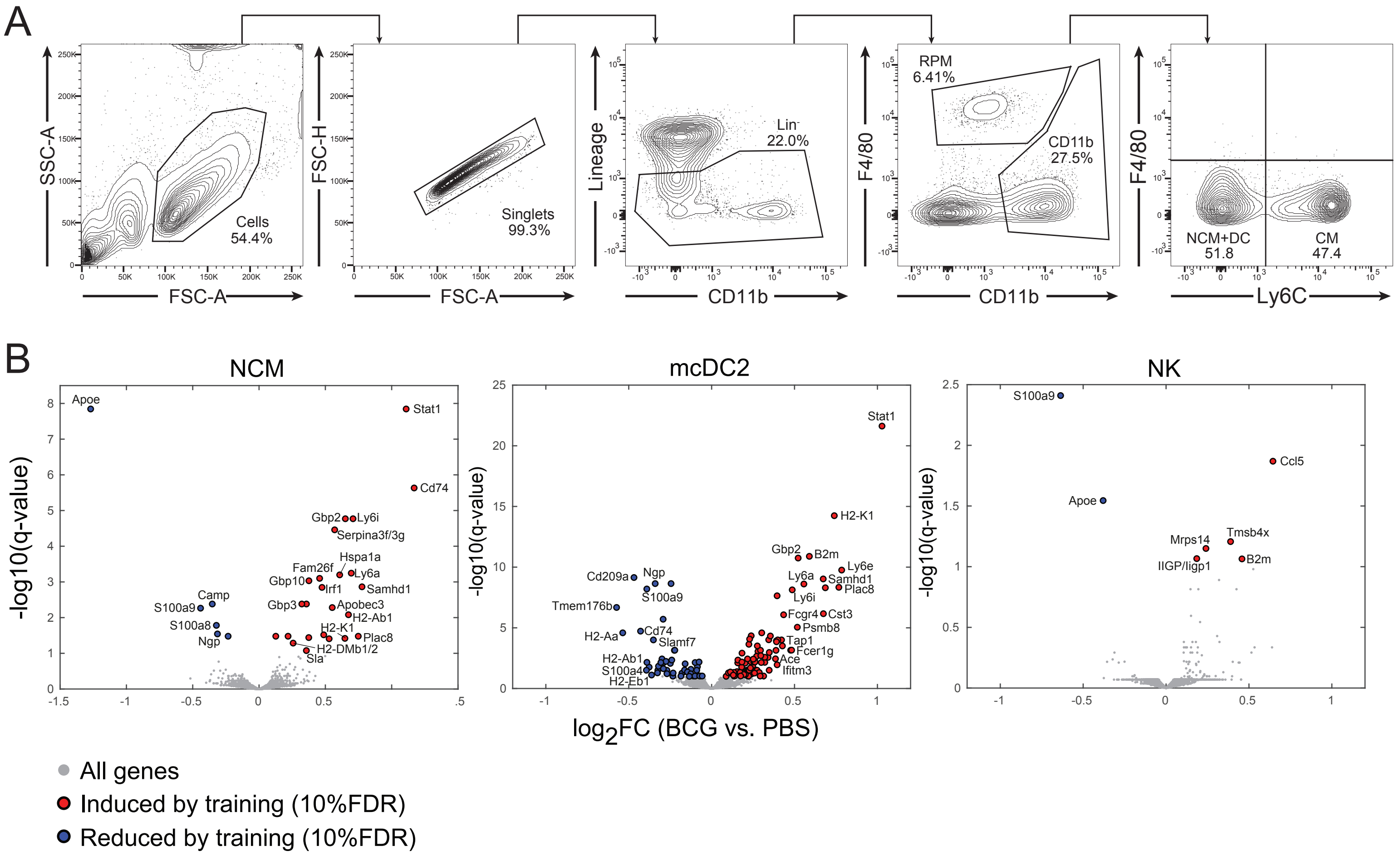

### Supplement Figure 2

#### Figure supplement 2

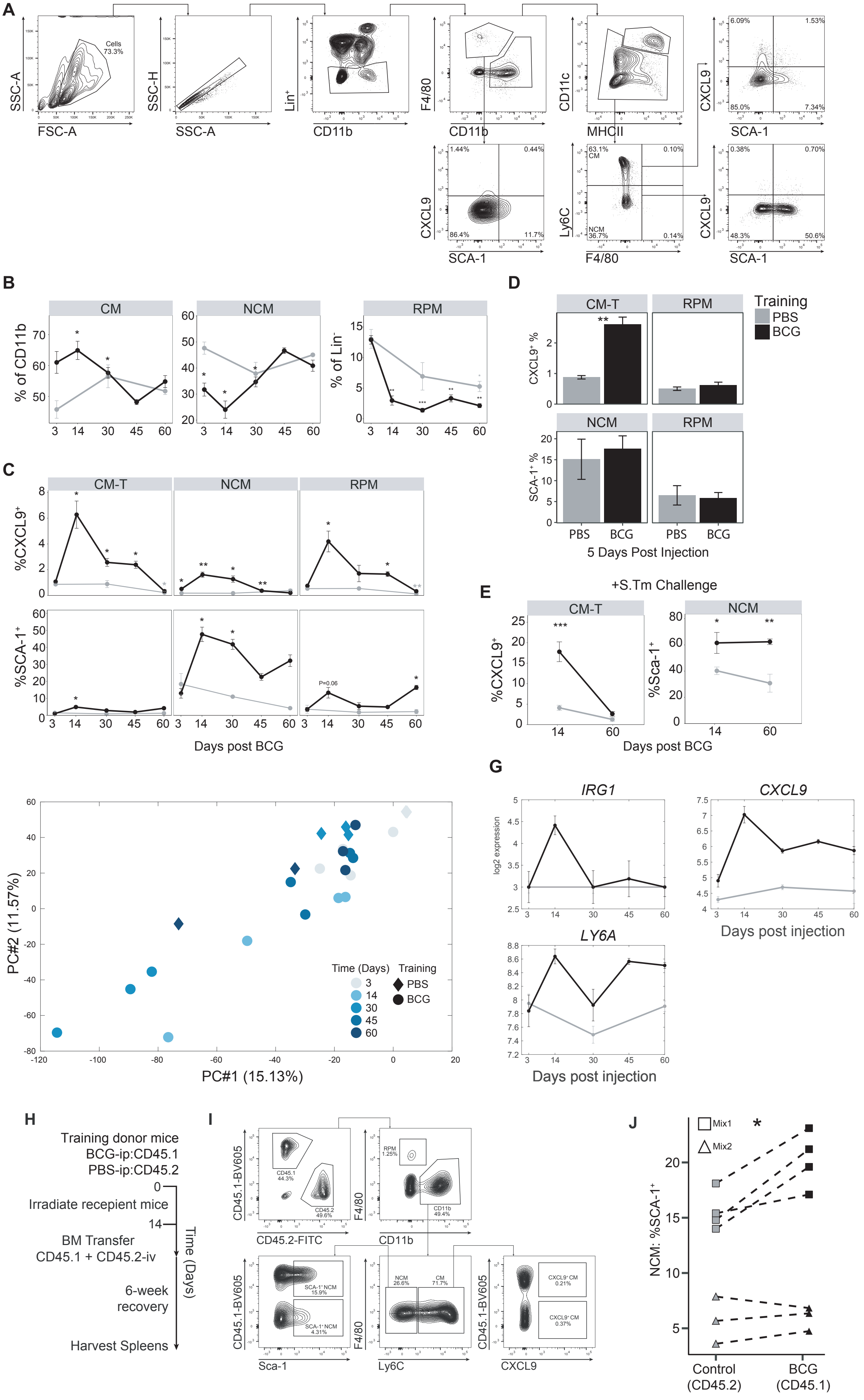

### Supplement Figure 3

##### Figure supplement 3

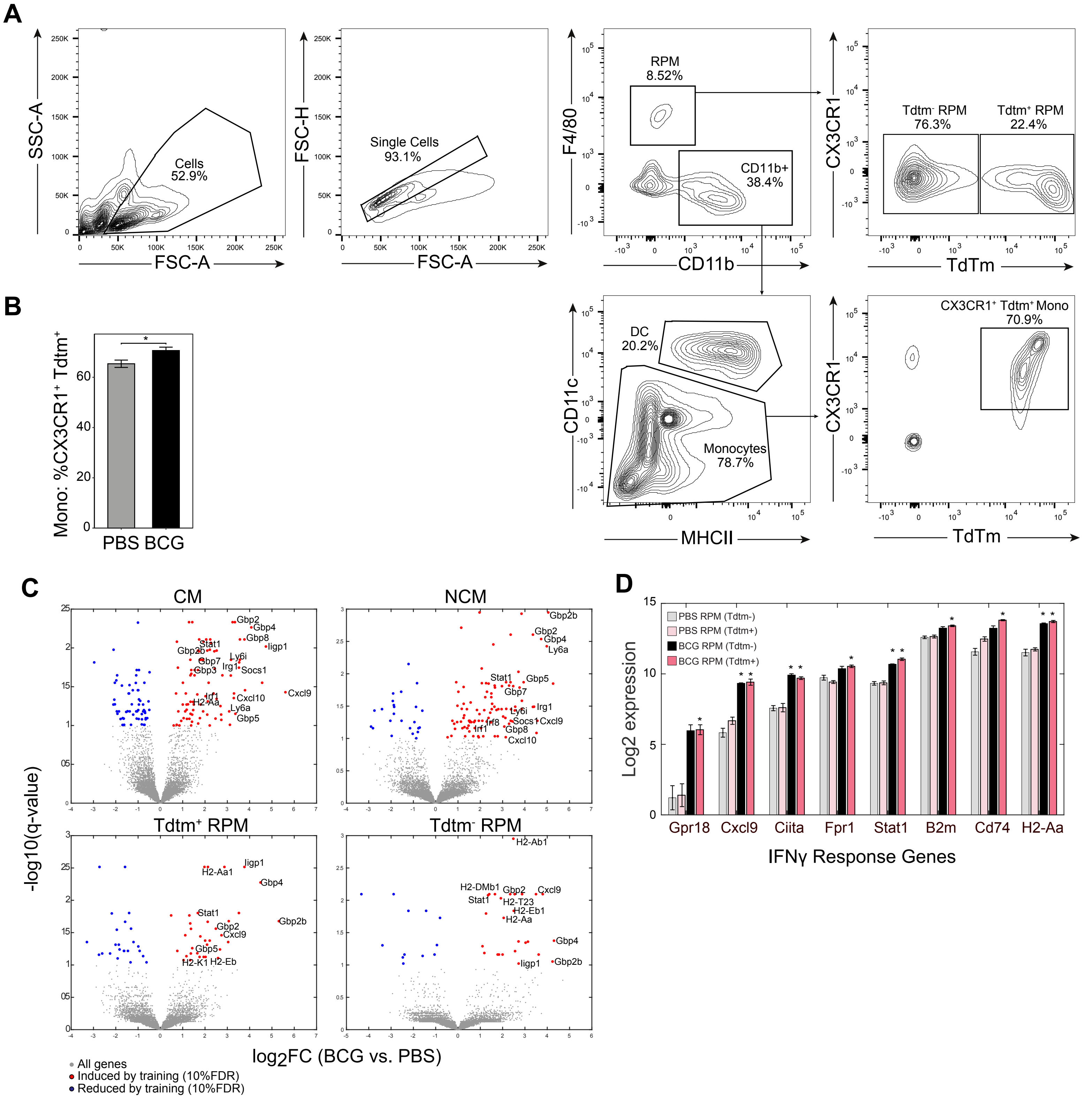

### Supplement Figure 4

**Figure supplement 4**

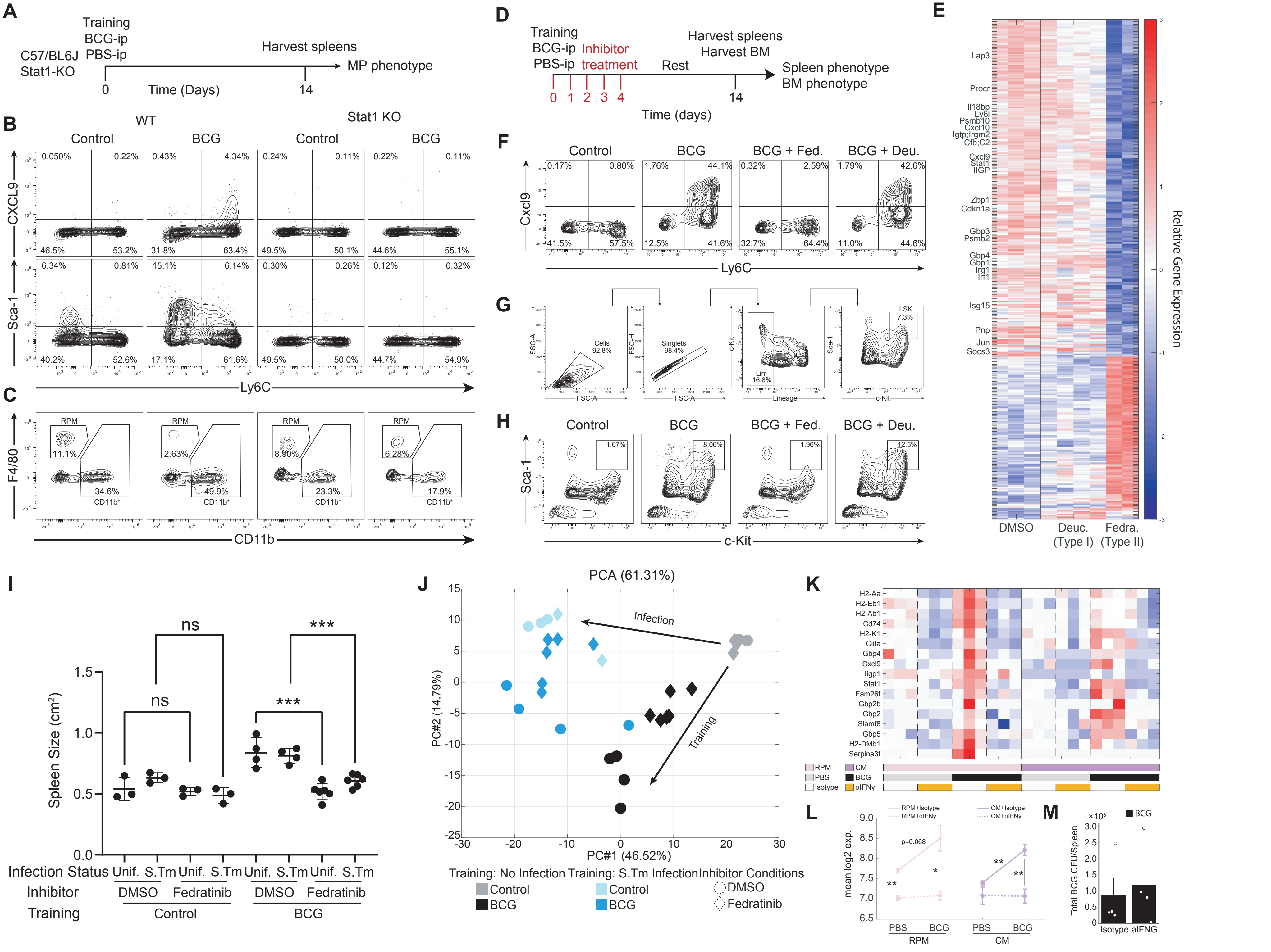
